# Endosomal MrGPRX1 signaling sensitizes TRPV1 to enhance itch

**DOI:** 10.1101/2025.11.12.688053

**Authors:** Paz Duran, Jeffri S. Retamal, Marcella de Amorim Ferreira, Kai Trevett, Dane D. Jensen

## Abstract

G protein-coupled receptors (GPCRs) and TRPV (transient receptor potential vanilloid) channels are crucial for signal transduction in physiological processes, including neurotransmission, pain, and itch. Downstream effectors of GPCR signaling can directly stimulate TRPV channels or enhance their sensitivity to stimuli, a process known as TRPV sensitization. Traditionally, GPCRs are activated at the cell surface by extracellular agonists, triggering signaling cascades. Recent evidence suggests GPCRs continue to signal from intracellular organelles. The human Mas-related G-protein coupled receptor X1 (MrGPRX1) is a GPCR expressed in primary sensory neurons involved in nociception and pruritus. Recent studies demonstrated how intracellular GPCR signaling regulates neuronal activity. However, there is no evidence characterizing MrGPRX1 trafficking or intracellular signaling. Herein, we characterized MrGPRX1 signaling within the endosomal network and its role in sensitizing TRPV1 channels to enhance itch signaling. Utilizing subcellular targeted biosensors, we demonstrated MrGPRX1 can traffic and signal from endosomes. Immunofluorescence analysis showed that MrGPRX1 internalizes following BAM8-22 stimulation. BRET assays revealed that MrGPRX1 activation induces Gα_q_ and β-arrestin-1 recruitment to the plasma membrane and early endosomes. Inhibition of dynamin or clathrin blocked BAM8-22-induced MrGPRX1 endocytosis and decreased nuclear extracellular signal-regulated kinase (ERK) signaling. Calcium signaling confirmed that MrGPRX1-mediated TRPV1 sensitization is mediated by protein kinase C and ERK activation. Our findings reveal a novel role for MrGPRX1 endosomal signaling in TRPV1 sensitization.

Understanding the mechanisms of MrGPRX1 signaling offers valuable insights into differentiating between pain and itch pathways, aiding in the development of targeted therapies for chronic pain and persistent itch.

## Main Text Introduction

Pruritus (itch) is a significant unresolved clinical issue, described as an unpleasant sensation that provokes the desire to scratch. This conserved mammalian response serves as a physiological self-protective mechanism, physically removing foreign objects or irritants from the skin. However, chronic or intense acute itch can lead to great discomfort when not manage properly (1). Itch affects one in four adults, severely diminishing the quality of life, increasing stress levels and making itch one of the most debilitating conditions in patient care (2, 3). Despite impacting one in four adults, there are few therapies to manage chronic itch as the molecular, cellular, and neural circuit mechanisms underlying this condition are not yet fully understood. Thus, it is essential to characterize the molecular and cellular mechanisms that regulate itch in order to develop targeted therapies and improve patient outcomes.

Itch irritants on the skin are detected by peripheral sensory neurons expressing specific receptors and ion channels that respond to diverse thermal, mechanical, and chemical noxious stimuli. Two key classes of itch sensing receptors are G protein-coupled receptors (GPCRs) and transient receptor potential (TRPs) ion channels. GPCRs are essential pruritogen detectors, detecting most itch inducing molecules controlling both histamine dependent and independent itch pathways(4, 5). Similarly, TRP channels like the transient receptor potential vanilloid 1 (TRPV1) play a large role in regulating itch sensation(6). A greater understating of the interplay between GPCRs and TRP channels is necessary as GPCRs are known to be modulate TRP channels, altering sensory neuron activity and gating between itch and pain sensation (7–11).

TRPs are a group of non-selective cation channels expressed in nociceptors that allow ions to pass through the plasma membrane(12). These channels are crucial for detecting and responding to noxious stimuli; therefore, they play critical roles in pain, inflammation, and itch (6, 13–16). TRP channels are a major downstream target for GPCR signaling, and the GPCR-TRP axis is vital for pain, neurogenic inflammation, edema and itch (16–19). Second messenger pathways generated from activated GPCRs can alter TRP channel activity and increase the expression of TRP channels at the cell surface. Additionally, GPCR activation can directly activate TRP channels (GPCR-TRP channel coupling) or enhance TRP responsiveness to TRP activators, a process known as TRP sensitization (16).

Mas-related G-protein-coupled-receptors (MrGPRs) are a family of orphan GPCRs that play important roles in various somatosensory functions, are key receptors in itch sensation and also have a role in nociception. MrGPRs are expressed in peripheral sensory neurons, dorsal root ganglion (DRG), and trigeminal ganglion (TG) (20–22), and have been associated with histamine dependent and histamine-independent itch signaling pathways (4, 23, 24). Within the MrGPR family, the primate-specific subfamily X, comprised of 4 receptors, has emerged as promising pharmacological targets due to their roles in pain, inflammation, neuroimmune diseases, allergies and itch (4, 16, 25). MrGPRX1, in humans, has been identified as a receptor for pruritogens such as the antimalarial drug chloroquine (CQ). Furthermore, the endogenous pruritogen fragment Bovine Adrenal Medulla peptide (BAM) 8-22, a proenkephalin A gene product, exhibits the highest specificity for human MrGPRX1 (26). BAM8-22 binds and activates MrGPRX1 found on primary sensory neurons to transmit itch signals from the periphery to the central nervous system. MrGPRX1 is a Gα_q_ coupled GPCR and once activated at the cell surface by extracellular agonists, facilitates intracellular Ca^2+^ release and ion channel activation(27, 28). Following activation, most GPCRs are endocytosed to the endosomal network. Previous studies have demonstrated that GPCRs can exhibit sustained compartmentalized signaling from intracellular organelles, and this prolonged signaling is significant in neuronal activation associated with conditions like chronic pain (29, 30). Given the established role of MrGPRX1 in mediating itch, we hypothesize that MrGPRX1 can signal from intracellular compartments, and that endosomal signaling of MrGPRX1 is crucial for sensitizing TRPV1 channels, leading to enhanced itch sensation.

## Results

### BAM8-22 Stimulates MrGPRX1 Endocytosis and Trafficking to Endosomes

MrGPRX1 trafficking after activation was followed by specific labeling with the mApple red fluorescent protein genetically fused to the intracellular C-terminus of the receptor. We transiently expressed MrGPRX1-mApple in HEK293 cells and tracked MrGRPX1-mApple internalization using confocal microscopy. In vehicle-treated cells, fluorescent MrGPRX1-mApple was principally localized at the plasma membrane (**Fig. 1A**; Control; arrowheads). After BAM8-22 treatment (1 μM, 15 min), MrGPRX1-mApple was depleted from the plasma membrane and accumulated in intracellular vesicles (**Fig. 1A**; BAM8-22; arrows). Vesicles were co-stained with an antibody to the early endosome antigen 1 (EEA1), a marker for early endosomes(30, 31). Thus, BAM8-22 stimulation induced MrGPRX1-mApple trafficking from the plasma membrane to early endosomes. Inhibition of dynamin mediated endocytosis was accomplished by the overexpression of the dominant negative mutant of dynamin (DynK44A)(32). Co-expression of DynK44A inhibited agonist-stimulated MrGPRX1-mApple internalization (**Fig. 1A**; DynK44A; arrowheads).

**Figure 1.**
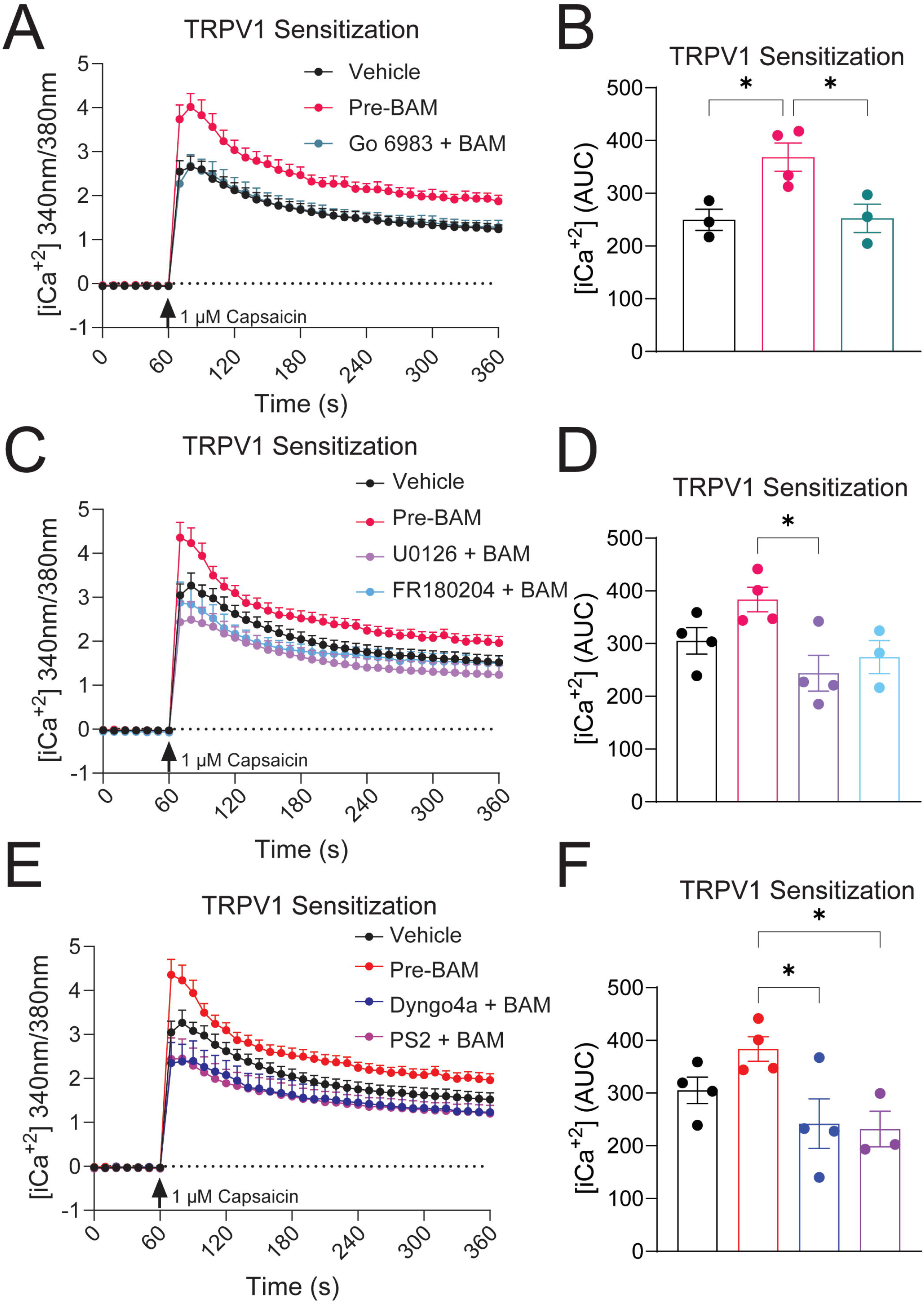
MrGPRX1 endocytosis can be blocked by dynamin inhibition. (A) Plasma membrane bound MrGPRX1-mApple is endocytosed to early endosomes (EEA1 label) 15 min following the addition of BAM8-22 in HEK293 cells. MrGPRX1 endocytosis is blocked by the expression of dominant negative mutant of dynamin (DynK44A). Arrowheads, cell-surface; arrows, endosomal MrGPRX1-mApple. Scale 10 µm. Representative images, n=3 independent experiments. BAM8-22-induced (1 µM) trafficking of MrGPRX1-Rluc8 from the (B) plasma membrane (CAAX-Venus) to (C) early endosomes (Rab5a-Venus) in HEK293 cells is blocked by DynK44A expression. Area under the curve (AUC) represents quantification of change in BRET over time. n≥5, * P≤ 0.05, ** P≤ 0.01. Unpaired t-test.

Bioluminescence resonance energy transfer (BRET) assays were used to quantify the trafficking and colocalization of MrGPRX1 in the endosomal compartment. HEK293 cells were transfected to express MrGPRX1 tagged with the Renilla Luciferase (MrGPRX1-RLuc8) and various subcellular markers to identify the plasma membrane (CAAX-Venus), or early endosomes (Rab5a-Venus) coupled to the BRET acceptor Venus(30). MrGPRX1-RLuc8 expressing cells were stimulated with BAM8-22 (1 μM) and the changes in BRET signal were measured. BAM8-22 activation of MrGPRX1-RLuc8 decreased the BRET signal between MrGPRX1-RLuc8 and the plasma membrane marker (**Fig. 1B**; CAAX-Venus), and increased BRET signal between MrGPRX1-RLuc8 and the early endosomes marker (**Fig. 1C**; Rab5a-Venus) denoting receptor internalization and trafficking away from the plasma membrane to early endosomes. Additionally, BAM8-22 stimulation resulted in an increase in BRET signal between MrGPRX1-RLuc8 and the late endosome marker (**Supporting** **Figure 1A and 1B**; Rab7a-Venus), and recycling endosomes (**Supporting** **Figure 1C and 1D**; Rab11a-Venus). Furthermore, co-expression of DynK44A significantly decreased the agonist-stimulated change in BRET (**Fig. 1B and 1C**). These results demonstrate that MrGPRX1 traffics from the plasma membrane to the endosomal pathway after BAM8-22 stimulation in a dynamin dependent process.

### MrGPRX1 Endocytosis Mediates Signaling in Subcellular Compartments

To assess if MrGPRX1 can persist in an active state and recruit β-arrestin isoforms and G proteins to early endosomes, we used nanobit-BRET (nbBRET) assays. The nbBRET assays were designed to enable the measurement of the interaction between three proteins by splitting the NanoLuc luciferase (NLuc) into 2 peptides, a small 13 amino acid peptide known as NanoBiT fragment (NP) and a larger fragment known as LgBiT (33, 34). When these two peptides are in close proximity (<10 nm), they form a functional NLuc luciferase that can emit luminescence in the presence of its substrate furimazine. This approach allows us to measure the ability of MrGPRX1 to recruit effector proteins to different subcellular compartments. For this, we transfected HEK293 cells with a C-terminal tagged MrGPRX1 with NP (MrGPRX1-NP) and either a plasma membrane marker (CAAX-LgBiT) or an early endosome marker (FYVE-LgBiT), and different effector proteins coupled to the BRET acceptor Venus (**Fig. 2A**).

**Figure 2.**
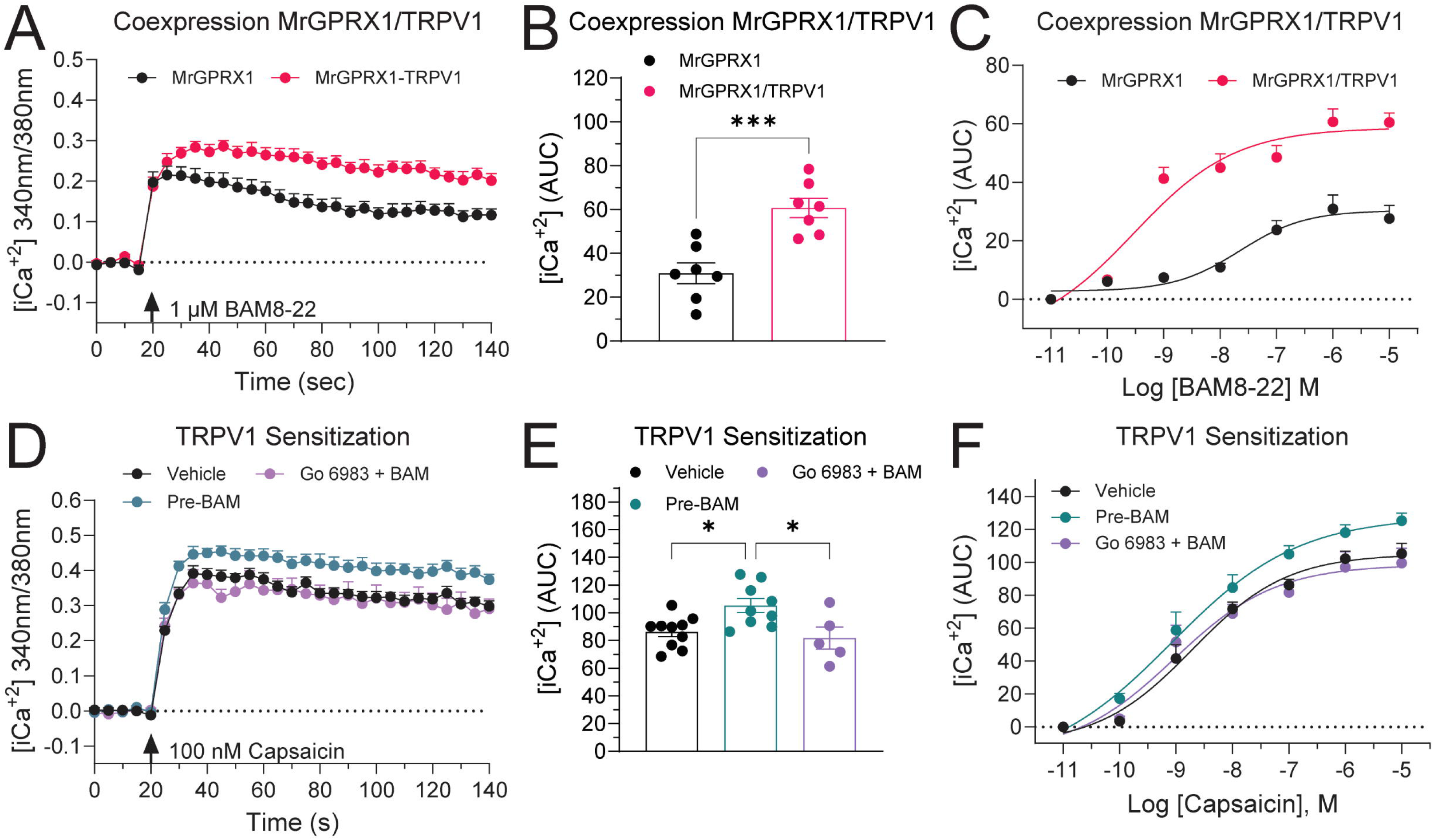
MrGPRX1 recruits miniGα_q_ (mGα_q_) and β-arrestin 1 (βArr1) to plasma membrane and early endosomes. (A) Illustration of NanoBit-BRET (nbBRET) assays. nbBRET uses a split luciferase (NanoBit or NP and LgBiT) that when in close proximity form a functional nanoluciferase to detect BRET between three different proteins; receptor (MrGPRX1), effector (mGα_s/q_ or βarr1), and proteins resident in subcellular compartments. Plasma membrane (PM). BAM8-22 induced recruitment of Venus-mGα_q_ to (B) the plasma membrane (CAAX-LgBit) and (E) early endosomes (FYVE-LgBiT) in HEK293 cells. BAM8-22 induced recruitment of Venus-βArr1 to (C) the plasma membrane (CAAX-LgBit) and (F) early endosomes (FYVE-LgBiT) in HEK293 cells. (D,G) Area under the curve (AUC) represents quantification of change in nbBRET over time. n≥4

We first evaluated the recruitment of G proteins to MrGPRX1-NP. For this, we used miniGα proteins (mGα) which are N-terminally truncated Gα isoforms that can bind to active conformations of GPCRs (35, 36). It has been reported that MrGPRX1 recruits and associates to Gα_q_ (23, 27, 37). Consistent with this, stimulation with BAM8-22 (1 μM) caused an increase in nbBRET signal at the plasma membrane between MrGPRX1-NP, CAAX-LgBiT and Venus-mGα_q_ but not Venus-mGα_s_ (**Fig. 2B and 2D**). When the endosomal marker FYVE-LgBiT was transfected, an increase in nbBRET in response to BAM8-22 (1 μM) was detected between MrGPRX1-NP and Venus-mGα_q_ but not Venus-mGα_s_ (**Fig. 2E and 2G**), showing that Gα_q_ proteins can be recruited to endosomal compartments containing the active MrGPRX1. Additionally, BAM8-22 (1 µM) activation induces the recruitment and association of β-arrestin 1-Venus but not β-arrestin 2-Venus with MrGPRX1-NP at the plasma membrane (**Fig. 2C and 2D**). A similar pattern was observed at early endosomes, where BAM8-22 (1 μM) induced an increase in nbBRET between MrGPRX1-NP, FYVE-LgBiT and β-arrestin 1-Venus but not β-arrestin 2-Venus (**Fig. 2F and 2G**). Taken together, these results show that following activation by BAM8-22, MrGPRX1 is bound by β-arrestin 1, undergoes dynamin-dependent endocytosis and traffics to early endosomes where activated MrGPRX1 can recruit effector proteins like mGα_q_.

Given the important involvement of ERK signaling in regulating crucial cellular responses, we investigated the contribution of MrGPRX1 endocytosis to ERK activation. This was monitored with high spatial and temporal resolution using Förster resonance energy transfer (FRET) based ERK biosensors known as EKAR biosensors. These contain a reversible substrate sequence separated by two fluorophores and can be targeted to different subcellular compartments (38, 39). We measured cytosolic (CytoEKAR) and nuclear (NucEKAR) ERK activity in HEK293 cells expressing human MrGPRX1. BAM8-22 (1 μM) induced robust and sustained ERK activation in the cytosol (**Fig. 3A-C**) and nucleus (**Fig. 3D-F**). Interestingly, pretreatment with the clathrin inhibitor Pitstop2 (PS2) significantly decreased BAM8-22-induced activation of both cytosolic (**Fig. 3A and 3C**) and nuclear (**Fig. 3D and 3F**) ERK, while the inactive version of Pitstop2 (PS2 Inact) had no significant impact on ERK signaling.

**Figure 3.**
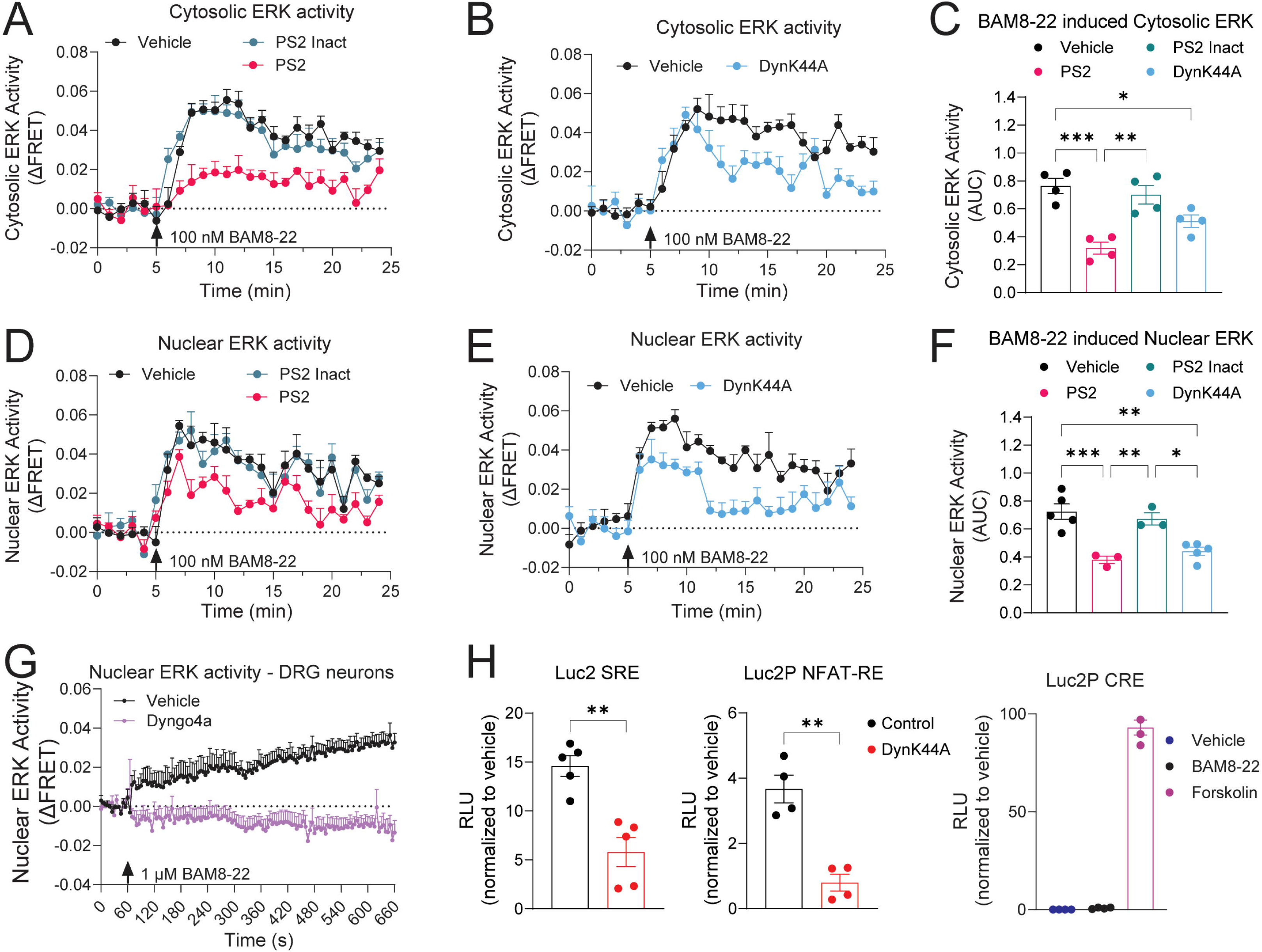
Characterization of the compartmentalized subcellular signaling of MrGPRX1. ERK phosphorylation was determined using EKAR FRET biosensors. BAM8-22 induced activation of cytosolic (A-C) and nuclear (D-F) ERK in HEK293 cells. Pretreatment with endocytic inhibitor PS2 (A, D) or expression of DynK44A (B, E) attenuate ERK activation. (C, F) Area under the curve (AUC) represents quantification of ERK activation over time. n≥3, * P≤ 0.05, ** P≤ 0.01, *** P≤ 0.001. One-Way ANOVA, Tukey’s multiple comparisons. (G) Pretreatment with the dynamin inhibitor Dyngo4a reduced nuclear ERK activation induced by BAM8-22 in mouse DRG neurons. (H) Luciferase transcriptional assays of HEK293 cells expressing MrGPRX1 and Luc2-SRE, Luc2-NFAT-RE, or Luc2-CRE in response to BAM8-22 (1 µM, 4 h). Expression of DynK44A attenuated MrGPRX1-induced transcription. Relative luminescence units (RLU). n≥4, ** P≤ 0.01. Unpaired t-test.

Furthermore, in cells expressing DynK44A, ERK activity was significantly reduced in both compartments compared to the control condition (**Fig. 3B, C, E, F**). Given that MrGPRX1 is mainly expressed on primary sensory neurons, we used an adeno-associated virus (AAV-NucEKAR) to express the ERK biosensor specifically in the nucleus of mouse dorsal root ganglion (DRG) neurons (**Supporting** **Figure 2**). Similar to our results in HEK293 cells, treatment with BAM8-22 (1 μM) produced a sustained nuclear ERK response in mouse DRG neurons. This activity was prevented by preincubation with the dynamin inhibitor, Dyngo4a (10 μM) (**Figure 3G**). Thus, these results highlight the necessity of endosomal trafficking and signaling of MrGPRX1 in driving a complete cellular response to MrGPRX1 activation.

Nuclear ERK activation is a crucial step in the regulation of immediate early response genes that guide alterations in neuronal plasticity and neuronal networks (40, 41). We hypothesized that the activation of nuclear ERK via the endosomal signaling of MrGPRX1 would alter gene expression patterns that alter cellular responses. To further investigate the role of endosomal MrGRPRX1 on gene expression, we used transcriptional luciferase reporter assays(42). HEK293 cells were co-transfected with MrGPRX1 and luciferase reporter vectors. These vectors contain the luciferase gene (Luc2P) driven by promoters featuring specific response elements (REs). BAM8-22 (1 μM) induced transcriptional activity for elements responsive to both ERK signaling pathway (**Figure 3H**; Luc2P-serum response element, SRE), and Gα_q_ activation (**Figure 3H**; Luc2P Nuclear Factor of Activated T-cells response element, NFAT-RE). Moreover, the co-expression of DynK44A significantly decreased these transcriptional activities (**Figure 3H**). Consistent with previous reports, BAM8-22 failed to induce activity in response to cAMP (**Figure 3H**; Luc2P cAMP response element, CRE) compared to Forskolin (10 μM) used as a positive control (**Figure 3H**). These findings, together with the evidence of nuclear ERK signaling, establish MrGPRX1’s endocytosis and subsequent endosomal signaling as key requirements to generate the full cellular response to BAM8-22.

### MrGPRX1 Sensitization of TRPV1 Channels is Regulated by PKC

TRPV1 channels play crucial roles in the itch transmission (15). Studies show that MrGPRX1 can sensitize TRPV1 channels via Gα_q_-induced activation of protein kinase C (PKC) (43). To corroborate this, we measured changes in intracellular Ca^2+^ levels ([iCa^2+^]) using calcium imaging. BAM8-22 produced an increase in [iCa^2+^] in HEK293 cells expressing MrGPRX1. This calcium response was significantly enhanced by the co-expression of TRPV1 channels (**Figure 4A-C**), confirming a functional coupling between MrGPRX1 and TRPV1. Furthermore, as expected, pre-stimulation with BAM8-22 significantly potentiated capsaicin-induced [iCa^2+^]. This effect was prevented by the pre-treatment with the PKC inhibitor Go 6983 (100 nM) (**Figure 4D-F**). Thus, BAM8-22 mediated activation of MrGPRX1 sensitized TRPV1 channels in a PKC dependent manner.

**Figure 4.**
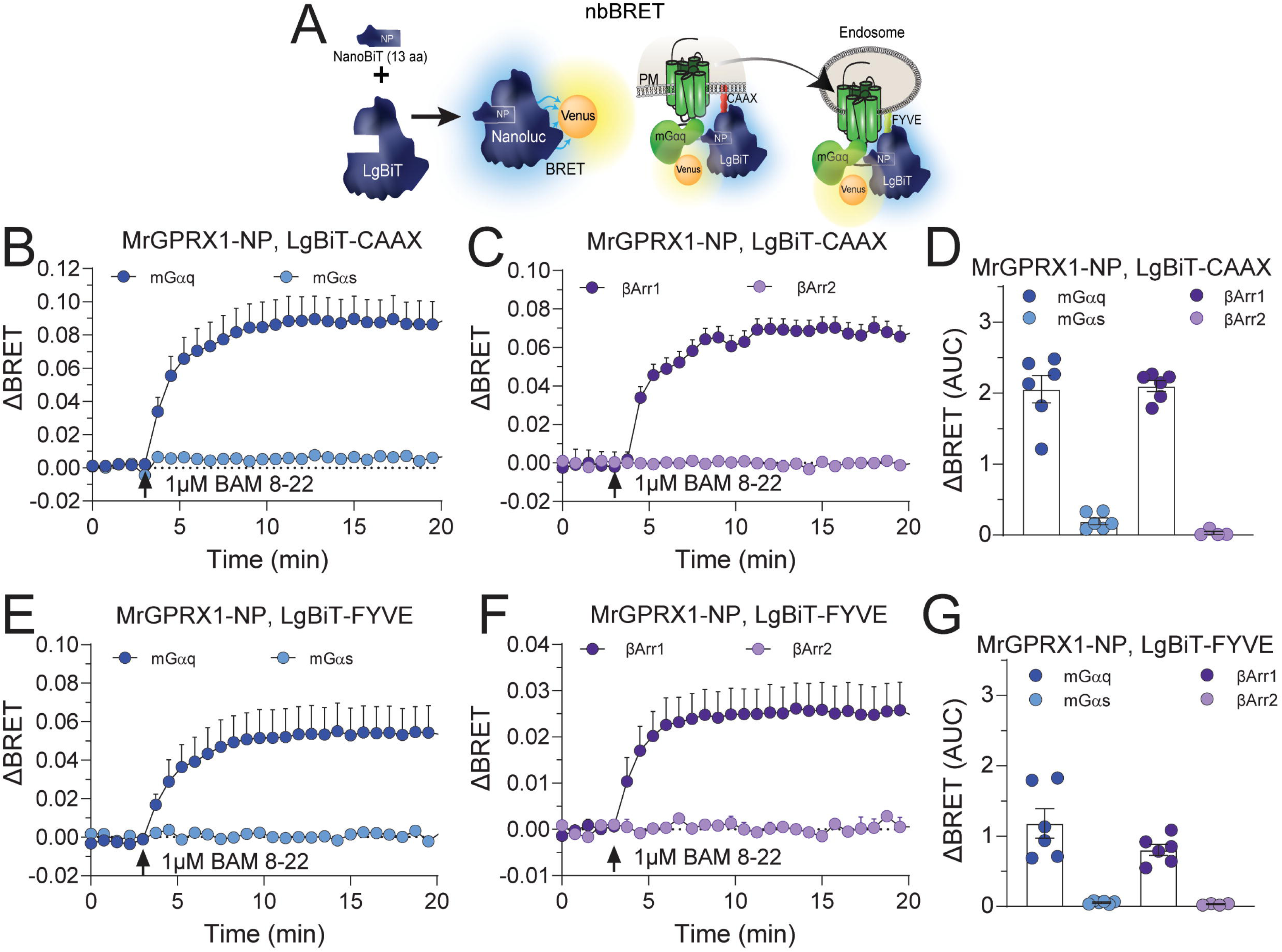
MrGPRX1 modulates TRPV1 channels. (A) BAM8-22-evoked Ca^2+^ signaling in HEK293 cells expressing MrGPRX1 alone or with TRPV1 channels. Area under the curve (AUC) represents quantification of intracellular Ca^2+^ change over time (B) and in a dose dependent manner (C). n=7, *** P≤ 0.001. Unpaired t-test. (D) Capsaicin-evoked Ca^2+^ influx in HEK293 cells expressing MrGPRX1 and TRPV1. MrGPRX1-mediated TRPV1 sensitization is decreased by pre-treatment with PKC inhibitor Go 6983. Area under the curve (AUC) represents quantification of intracellular Ca^2+^ influx over time (E) and in a dose dependent manner (F). n≥5, * P≤ 0.05. One-Way ANOVA, Tukey multiple comparisons.

### MrGPRX1 Sensitization of TRPV1 Channels in DRG neurons

To determine if MrGPRX1 also sensitizes TRPV1 channels in primary sensory neurons, we first confirmed their expression and co-localization in mouse DRG neurons. Although the individual expression of MrGPRX1 and TRPV1 in these neurons is well documented, a precise characterization of their co-localization has not been previously reported. For this, we used RNAScope® in situ hybridization. *Mrgprx1* (MrGPRX1) and *Trpv1* (TRPV1) mRNAs were detected in the same DRG neurons, which were identified by NeuN immunostaining (**Figure 5A**). *Mrgprx1* was detected in ∼18% and *Trpv1* in ∼26% of the neurons (**Fig 5B**). *Mrgprx1* and *Trpv1* were co-expressed in approximately 8% of total DRG neurons (**Fig 5B**).

**Figure 5.**
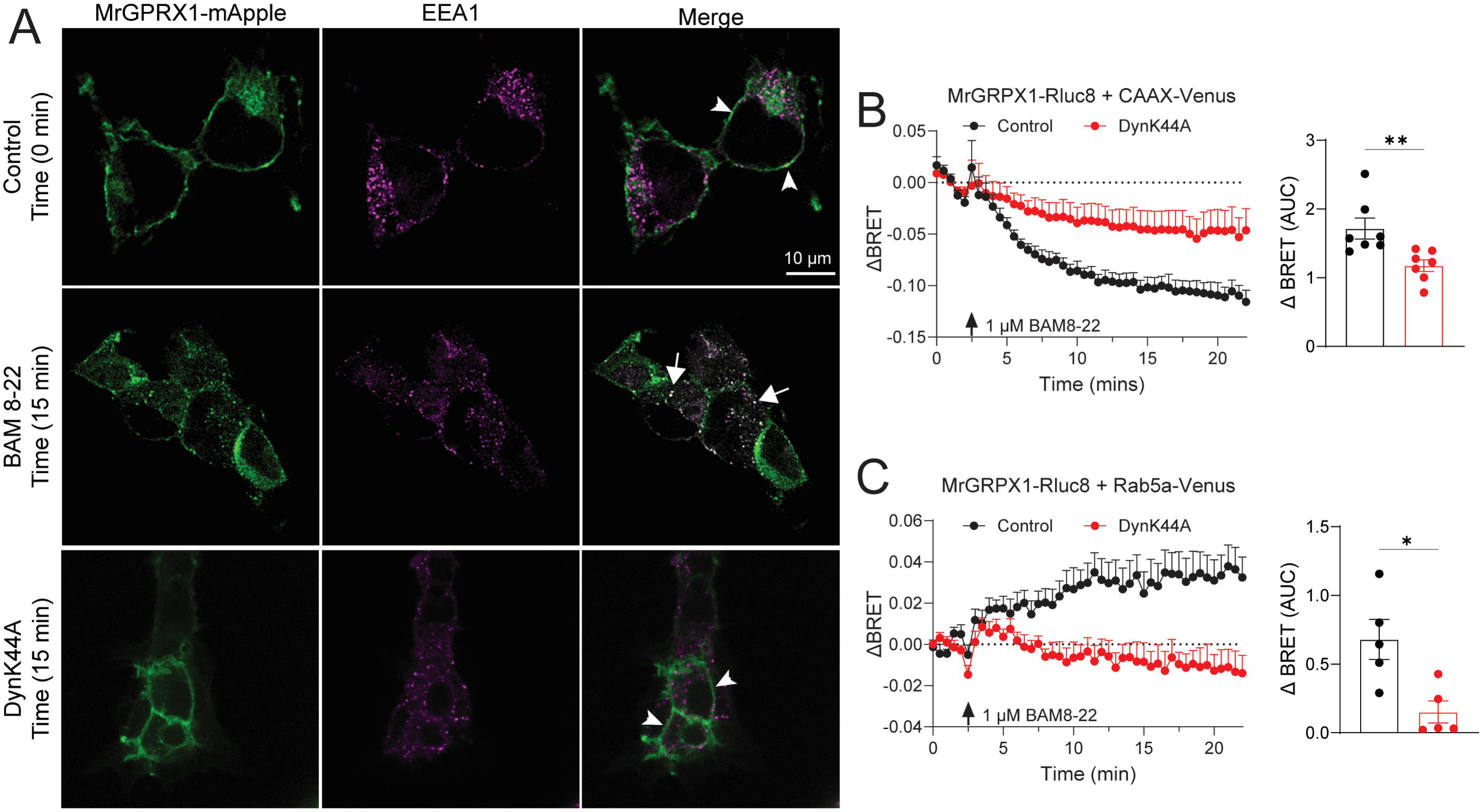
Characterization of MrGPRX1 and TRPV1 expression in mouse dorsal root ganglion (DRG) neurons. (A) Representative RNAScope® image of the detection of MrGPRX1 and TRPV1 mRNA in mouse DRG neurons identified by NeuN immunofluorescence. Arrows represent MrGPRX1 positive neurons, and arrowheads represent TRPV1 positive neurons. Scale, 100 µm, and 50 µm in inset. (B) Quantified percentages of MrGPRX1 positive neurons, TRPV1 positive neurons, and co-expression of MrgprX1 and TRPV1 in DRG neurons. N= 4 mice.

To further validate MrGPRX1 sensitization of TRPV1 channels, we performed live cell calcium imaging using isolated mouse DRG neurons. Results showed that pre-treatment with BAM8-22 (1 µM) significantly increased capsaicin-induced calcium influx through TRPV1 (**Figure 6**). Similar to what we observed on HEK293 cells, this increased Ca^2+^ influx was prevented by the pre-incubation with the PKC inhibitor Go 6983 (100 nM) (**Figure 6A and 6B**). These results confirm that PKC activation is essential for MrGPRX1 mediated sensitization of TRPV1 channels in DRG neurons.

**Figure 6.**
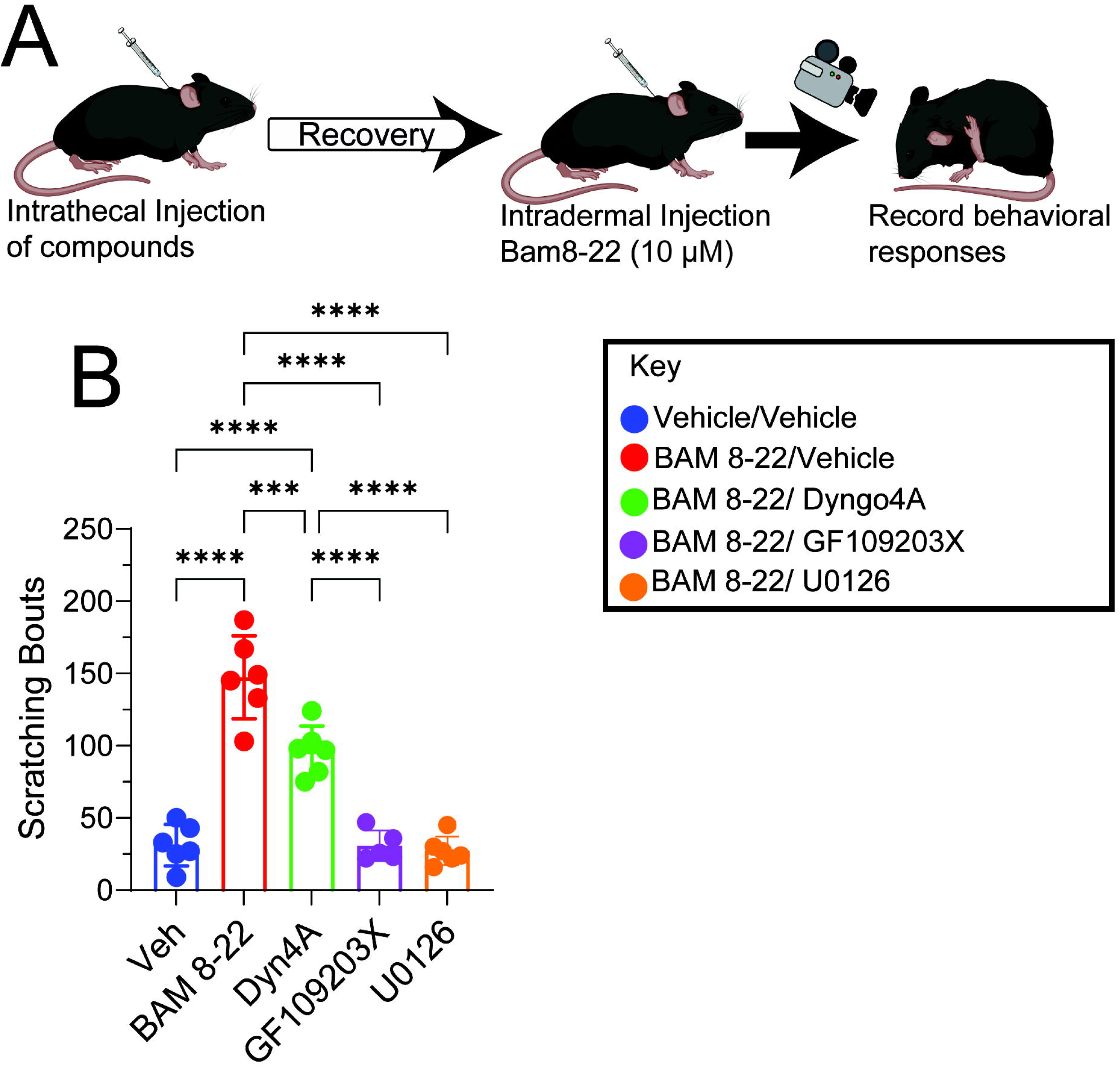
MrGPRX1 sensitizes TRPV1 channels in mouse DRG neurons. Capsaicin-evoked Ca^2+^ influx in mouse DRG neurons. BAM8-22-mediated TRPV1 sensitization is decreased by pre-treatment with PKC inhibitor Go 6983 (A-B), MEK inhibitor U0126 (C-D), ERK inhibitor FR 180204 (C-D), and endocytic inhibitors Dyngo4a or PS2 (E-F). Area under the curve (AUC) represents quantification of intracellular Ca^2+^ influx over time. n≥3 mice, * P≤ 0.05. One-Way ANOVA, Tukey multiple comparisons.

In addition, we investigated the role ERK signaling in TRPV1 sensitization in response to BAM8-22. For this, we pre-incubated the neurons with the selective mitogen-activated protein kinase (MEK) inhibitor U0126 (10 µM; 30 min) to inhibit the MEK/ERK signaling cascade. Calcium imaging results demonstrated that TRPV1 sensitization by BAM8-22 was significantly blocked by the pre-incubation with U0126 (**Figure 6C and 6D**). Furthermore, to confirm that ERK activity is regulating TRPV1 sensitization, we also tested the selective ERK inhibitor FR 180204 (30 µM; overnight). The results showed that ERK inhibition partially prevented the sensitization of TRPV1 channels (**Figure 6C and 6D**). These findings indicate that ERK activation, in conjunction with PKC activation, regulate BAM8-22 mediated sensitization of TRPV1 channels.

Next, we investigated the role of endosomal MrGPRX1 signaling in TRPV1 sensitization. For this, we pre-incubated mouse cultured neurons with the endocytic inhibitors Dyngo4a (10 μM) or PS2 (10 μM) and then performed calcium imaging. Both inhibitors effectively suppressed the BAM8-22 induced sensitization of TRPV1 channels (**Figure 6E and 6F**), validating our hypothesis that endosomal signaling of MrGPRX1 is crucial for the sensitization of TRPV1 channels in DRG neurons.

### Endosomal MrGPRX1 signaling mediates scratching behaviors

Calcium imaging results confirmed that BAM8-22 sensitization of TRPV1 is regulated by MrGPRX1 endosomal signaling and both MRGPRX1 and TRPV1 have been shown to control a scratching response in mice(6, 24). To evaluate the role of endosomal MrGPRX1 signaling in scratching behavior, we first injected intrathecally the endocytic inhibitor Dyngo4a or vehicle control. Mice were allowed to recover 30 min before we examined the scratching behavior in response to an intradermal injection of BAM8-22 (10 μM). Scratching behavior, grooming, distance traveled, and inactivity were monitored using the Behavioral Spectrometer (**Figure 7A**). BAM8-22 administration caused a significant increase in total scratching bouts (**Figure 7B**) compared to vehicle injected mice. Interestingly, pre-administration of Dyngo4a (5 μl/30 μM) resulted in a significant decrease in scratching bouts in response to BAM8-22 compared to vehicle-treated mice (**Figure 7B**). To evaluate whether this scratching response is mediated by PKC activity and MEK/ERK signaling, we administered PKC inhibitor GF109203X (0.25 µg/5 µL) or MEK inhibitor U0126 (0.5 µg/5 μL) intrathecally before measuring the scratching response to BAM8-22. Notably, pre-administration of both inhibitors suppressed the scratching response to BAM8-22 (**Figure 7B**).

**Figure 7.**
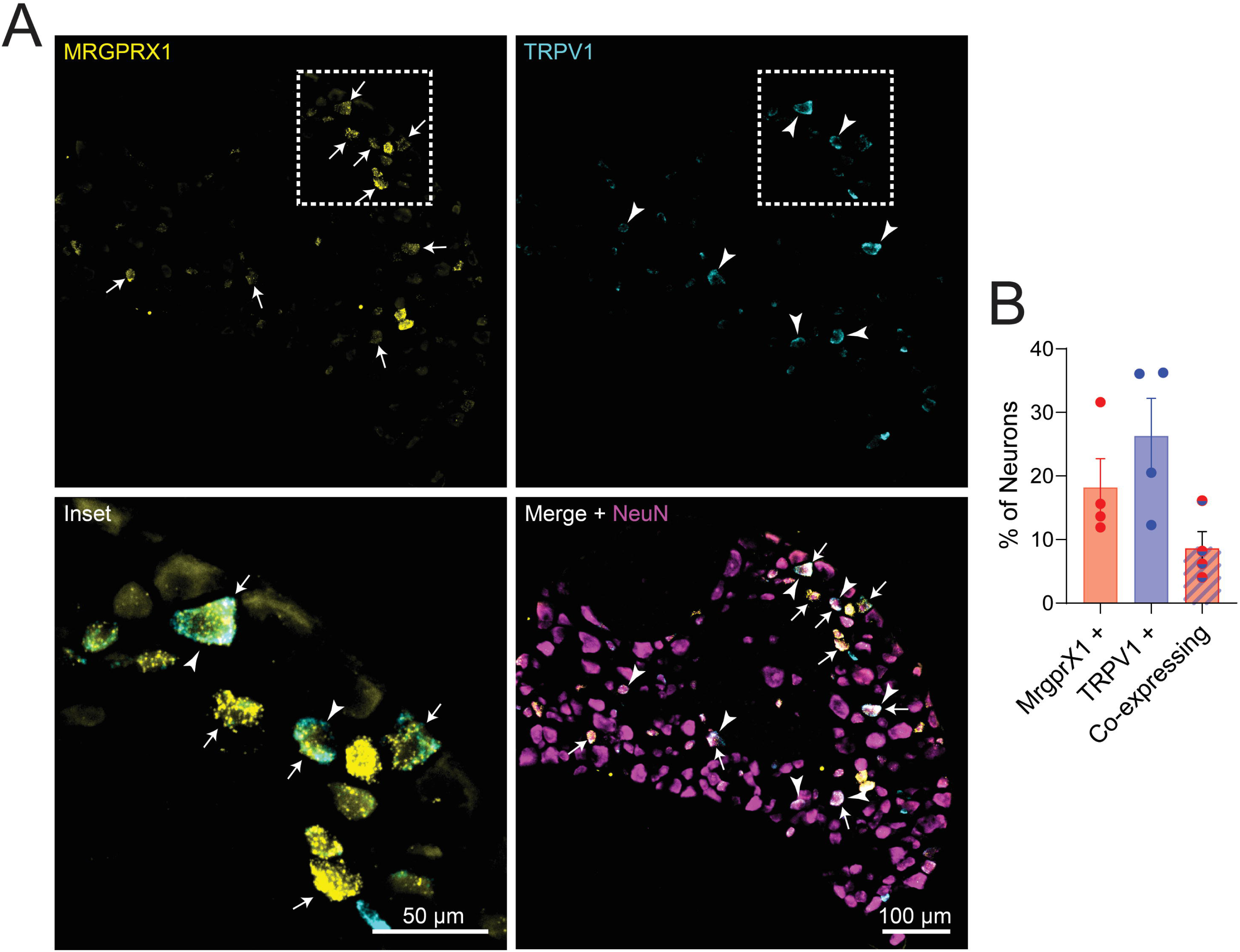
Effect of endocytic, MEK and PKC inhibitors in BAM8-22 induced scratching responses in mice. (A) Illustration of the experimental protocol. (B) Scratching bouts induced by vehicle or BAM8-22 intradermal injection 30 min after intrathecal administration of dynamin inhibitor Dyngo4A, MEK inhibitor U0126, or PKC inhibitor GF109203X. n>5 mice, *** P≤ 0.001, **** P≤ 0.0001. One-Way ANOVA, Tukey multiple comparisons.

Grooming, total distance traveled, and inactivity were not significantly altered by BAM8-22 administration. Furthermore, there was no significant change in grooming, distance traveled and inactivity in the Dyngo4a treated group (**Supporting** **Figure 3**). However, there was a significant decrease in the total grooming behavior for the group injected with the PKC inhibitor (GF109203X). Additionally, an increase in time inactive was observed in mice treated with MEK inhibitor (U0126) or PKC inhibitor (GF109203X) when compared with vehicle-treated mice (**Supporting** **Figure 3A and 3C**).

Collectively, these results confirm that internalization and endosomal signaling of MrGPRX1 in response to BAM8-22 is important to evoke a scratching response in mice, and this is likely mediated by PKC and ERK activation.

## Discussion and conclusion

This study characterizes a previously unknown signaling mechanism for the human MrGPRX1 receptor. For the first time, our results demonstrate that MrGPRX1, upon stimulation with BAM8-22, binds with β-arrestin-1 and undergoes clathrin and dynamin dependent endocytosis. Additionally, these studies show that MrGPRX1 is able to recruit β-arrestin-1 and Gα_q_ to the endosomal compartment where endosomal MrGPRX1 regulates both cytosolic and nuclear ERK activation. These data confirm the known PKC-dependent MrGPRX1-mediated TRPV1 sensitization and unveils a new mechanism whereby endosomal signaling of MrGPRX1 regulates TRPV1 sensitization through a MEK/ERK dependent pathway. Supporting these mechanistic *in vitro* findings, we found that BAM8-22-induced scratching behaviors in mice are significantly attenuated when either endocytosis of MrGPRX1 or PKC/ERK signaling is impaired.

Overall, these results characterize a new signaling mechanism in which BAM8-22-induced activation of MrGPRX1 drives endosomal signaling that regulates TRP sensitization and itch transmission in primary sensory neurons.

The link between prolonged intracellular GPCR signaling events and pathological states, such as chronic pain and itch, is being increasingly emphasized in new studies (30, 31, 44, 45). Our findings support this emerging paradigm by demonstrating that the MrGPRX1 receptor, implicated in pruritus, undergoes agonist-induced endocytosis and signals from endosomes. In contrast, a previous report suggested that the human MrGPRX1 receptor is resistant to agonist-induced endocytosis (46). This discrepancy is likely attributable to important methodological differences used to study receptor internalization. First, the previous study utilized expression vectors generated from the SNSR4 gene, which represents an earlier sequence version of MrGPRX1. Second, in that report, MrGPRX1 cell surface levels were detected using ELISA experiments where cells were immediately transferred to ice following stimulation with BAM8-22. Given that low temperatures are known to inhibit endocytic pathways (47), this reported lack of receptor endocytosis may be a methodological artifact. In contrast, our study employed several rigorous qualitative and quantitative techniques, including BRET and confocal microscopy, to confirm that MrGPRX1 undergoes endocytosis following activation.

Furthermore, the observations that BAM8-22-mediated activation of MrGPRX1 results in Gα_q_ and β-arrestin-1 recruitment to early endosomes and that MrGPRX1-induced transcription was decreased upon blocking endocytosis provide functional evidence that MrGPRX1 continues to signal from early endosomes, thereby directly linking MrGPRX1 to the formation of signalosomes and providing a mechanism for prolonged intracellular signaling.

Sensitization of peripheral sensory neurons is a key factor in the pathophysiology of chronic pruritus. This peripheral sensitization can lead to alloknesis and hyperknesis, which are common symptoms in patients with chronic pruritus conditions (48). Unlike transient pruritus, chronic pruritus relies on sustained signaling and sensitization of the sensory neurons. Our data demonstrate that MrGPRX1 endosomal signaling is one mechanism that leads to this sustained activity, including prolonged nuclear ERK activation, which is a known driver of neuronal plasticity and long-term sensitization (41, 49, 50). This finding suggests that the spatial location of MrGPRX1 signaling may determine the nature of the itch response. This mechanism offers a therapeutic opportunity through biased agonism: by selectively targeting the endosomal signaling of MrGPRX1 while leaving the surface signaling intact, we could potentially treat chronic itch while avoiding unnecessary side effects.

Our finding that MrGPRX1 signals from endosomes provides the structural basis for its role in regulating TRPV1 activity. The sensitivity to a ligand, and the magnitude and duration of TRP activation, can be augmented by functional interactions with GPCRs, often termed ‘coupling’. This coupling is generally bidirectional, with the functional interaction of a GPCR and an ionotropic channel like a TRP sometimes leading to an augmentation of GPCR signaling as well (16). GPCRs typically modulate TRP activity via two general mechanisms. The first involves Gα-mediated activation of phospholipase C (PLC) and phospholipase A2 (PLA2), which cleave fatty acids and generate endogenous TRP activators such as arachidonic acid (AA) and phosphatidylinositol 4,5-bisphosphate (PIP2). The second mechanism is through activation of PKC and PKA, serine/threonine kinases that phosphorylate the intracellular C-tail of TRP channels to enhance cell surface expression and activity (7, 8, 10, 16, 51–55). TRP channels such as TRPV1 and TRPA1 are widely implicated in transducing GPCR activation into membrane depolarization in nociceptive neurons (53, 56).

Consistent with this established framework, our calcium imaging results demonstrate a coupling between MrGPRX1 and TRPV1, showing that MrGPRX1 can modulate TRPV1 activity through activation of PKC. Importantly, RNAScope *in situ* hybridization analysis provided anatomical evidence for this interaction, demonstrating that the mRNA for MrGPRX1 and TRPV1 co-localizes in a subset of DRG neurons. Furthermore, our data show that MrGRPX1 endocytosis and ERK activation are also critical to this sensitization mechanism, pointing to a novel role for prolonged MrGPRX1 endosomal signaling in driving this pathway. However, a critical gap remains; the specific link between Gα_q_ and β-arrestin-1 recruitment to early endosomes, sustained ERK activation and the subsequent TRPV1 sensitization still needs to be studied. Given that PKC activation typically occurs at the plasma membrane (57), we hypothesized that the fast sensitization of TRPV1 channels due to PKC activation provides the initial acute itch response, while the endosomal signaling mediated by ERK activation provides the prolonged and sustained sensitization that is crucial for the pathophysiology of chronic pruritus.

MrGPRX1 is exclusively expressed in humans and lacks a single-gene mouse ortholog. While it exhibits some sequence homology to murine MrgprA receptors (58), functional homology is instead showed by other rodent receptors. Specifically, MrgprA3 and MrgprC11 are considered functional orthologs of MrGPRX1 simply because they share the same ligands, chloroquine (CQ) and BAM8-22, respectively (4). Although this species difference represents a significant limitation for direct translational research, studies using mouse models have nevertheless provided fundamental insights into the *in vivo* function of MrGPRX1 signaling pathway. Beyond species differences, other limitations of our study include the use of pharmacological inhibitors of endocytosis and important signaling pathways such as PKC and MEK. These inhibitors could have possible off-target actions, meaning the observed effects might be due to disrupted signaling of other receptors and are unrelated to MrGPRX1 signaling itself. Supporting this concern, our results showed that administration of PKC or MEK inhibitors had an effect on some non-evoked behaviors, and previous reports have shown a decrease in locomotor activity after PKC inhibition with tamoxifen (59). Further studies employing genetic manipulation methods, such as conditional knockouts and the utilization of a “humanized” MrGPRX1 mouse line, will be necessary to validate the specific role of MrGPRX1 and its downstream signaling molecules in these complex *in vivo* behaviors.

In conclusion, we have demonstrated that, after activation, the human MrGPRX1 receptor traffics to endosomes and transmit sustained signals that regulate TRPV1 sensitization to underlie itch transmission. The endosomal signaling of GPCRs presents new challenges for understanding how GPCR signaling is regulated and how endosomal signaling of GPCRs contributes to disease states like chronic pruritus, while also opening new targets for therapeutic development.

## Experimental Procedures

### Drugs and Reagents

BAM8-22, and GF 109203X (GFX) were purchased from Tocris Bioscience (Bristol, UK). DMEM, Hank’s balanced salt solution (HBSS), and ProLong™ Glass, were purchased from Thermofisher (Waltham, MA). Coelenterazine h was purchased from Nanolight Technology (Pinetop, AZ). Nano-Glo® Luciferase substrate furimazine from Promega (Madison, USA). Fura2AM was purchased from Cayman Chemicals (Ann Arbor, MI). Dyngo 4a was purchased from Abcam (Cambridge, UK). Dyngo4aInactive were from Dr. Adam McCluskey (University of Newcastle, AU). All other reagents were from Sigma (St. Louis, MO) unless otherwise specified.

### cDNAs

Constructs for human MrGPRX1 (MrGPRX1-mApple, MrGPRX1-NP) were purchased from Twist Bioscience (San Francisco, CA, USA). The following plasmids and FRET biosensors were purchased from Addgene: CytoEKAR Cerulean/Venus (Cytosolic ERK sensor, 18679), NucEKAR Cerulean/Venus (Nuclear ERK sensor, 18681), and Dynamin K44A (dominant negative dynamin-1, 34683). The pGL4.33[luc2P/SRE/Hygro], pGL4․30[luc2P⁄NFAT-RE⁄Hygro], and pGL4․29 [luc2P⁄CRE⁄Hygro] vectors were purchased from Promega (Madison, USA).

For BRET studies, human MrGPRX1 construct was modified with Renilla® luciferase 8 (Rluc8). Rab5a-Venus (early endosome), Rab7a-Venus (late endosomes), Rab11-Venus (recycling endosome), were from N. Lambert (Augusta University Medical College of Georgia, GA). βArr1-YFP and βArr2-YFP were from M. Carson, Duke University, Durnham, NC.

For nbBRET studies, MrGPRX1-NP was cloned using Gibson Assembly with cDNA encoding human MrGRPX1 with an N-terminal POMC signal sequence and Flag tag in a pcDNA5/FRT vector. The 13 amino acid natural peptide fragment of nanoluciferase (GVTGWRLCERILA, NP) was insert to the C-terminus of MrGPRX1 via a flexible linker (LRPLGSSGGG). Localization markers CAAX (plasma membrane) or FYVE (endosome) tagged on the N-terminus with HA, a short linker (GGSG) and the LgBiT tag were donated by the Alex Thompson lab (New York University). Venus-miniGα_q_, and Venus-miniGα_s_ were from N. Lambert.

### Cell Culture

Human Embryonic Kidney 293 (Thermofisher) were culture in Dulbecco’s modified Eagle’s medium (DMEM) supplemented with 10% normal fetal bovine serum (FBS) and 100 U/mL penicillin-streptomycin at 37 °C and 5% CO_2_. Human Embryonic Kidney 293 cell tetracycline-inducible system (T-Rex™293, Thermofisher) expressing TRPV1 (HEK-TRPV1) were culture in DMEM containing 10% tetracycline free FBS, 100 U/mL penicillin-streptomycin, 100 µg/ml Hygromycin B Gold (InvivoGen), and 5 µg/ml Blasticidin (InvivoGen) at 37 °C and 5% CO_2_. The expression of TRPV1 was induced overnight with 0.1 μg/mL tetracycline.

### Transfection of Cell lines

HEK293-Flp-In cells were transfected using polyethylenimine (PEI, Polysciences; 1:6 DNA:PEI) diluted in a 150 mM NaCl solution.

### MrGPRX1 internalization

HEK293 cells were transfected with MrGPRX1-mApple with or without Dynamin K44A and were plated on PDL coated 12 mm cover slips in a 24 well plate. Following transfection (48 h), cells were challenged with vehicle or BAM8-22 (1µM) for 15- or 30-min at 37 °C. Cells were washed with PBS and fixed with PFA (4% paraformaldehyde, in PBS, 20 min, on ice). Cells were blocked in 5% normal horse serum (NHS), 0.1% Saponin in PBS (1 h, RT), and incubated with rabbit anti-EEA1 (1:1000, # PA1-063A, Invitrogen) overnight, 4°C. Cells were washed and incubated with anti-rabbit Alexa Fluor® 488 (1:1000, 1h, RT; Invitrogen). Cover slips were washed 3X with PBS and mounted on glass slides with ProLong® Diamont Antifade (Thermo Fisher). Sections were imaged using a Leica SP8 confocal microscope with HCX PL APO 63x (NA 1.40) oil objectives (Leica-Microsystems). Images were processed using ImageJ and figures were made in Adobe Illustrator.

### BRET transfections

For BRET assays of MrGPRX1 translocation to different organelles, HEK293 cells were transfected in 10 cm dishes with MrGPRX1-Rluc8 (1 µg) and Venus-CAAX (3 µg, plasma membrane), Venus-Rab5a (3 µg, early endosomes), Venus-Rab7a (3 µg, late endosomes), or Venus-Rab11a (3 µg, recycling endosomes). To disrupt Dynamin, HEK293 cells were co-transfected with Dynamin K44A (1 µg).

To characterize the recruitment of G proteins to the plasma membrane or to endosomes, HEK293 cells were transfected with human MrGPRX1-NP (2 µg) and HA-LgBiT-CAAX (2 μg) or HA-LgBiT-FYVE (2 μg), and either Venus-mGα_q_ (1 μg), Venus-mGα_s_ (1 μg), YFP-βArr1 (1 μg) or YFP-βArr2 (1 μg).

### BRET measurements

Following transfection (48 h), HEK293 cells were washed with HBSS containing 10 mM HEPES at pH 7.4. In some experiments, cells were pre-incubated in HBSS containing Pitstop 2 or Pitstop 2-negative control (10 µM; Abcam), Dyngo4a (10 µM; Abcam) or Dyngo4a Inactive (10 µM), or vehicle for 30 min, 37°C. Prior to BRET measurements, substrate was added for eBRET (coelenterazine H, 2.5 μM, 10 min), or nbBRET (furimazine, 10 µM, 10 min). BRET was recorded for up to 25 min in a CLARIOstar Microplate reader (BMG Labtech, Cary NC) using BRET and nbBRET filters (donor filter: 460 ± 40 nm, acceptor filter: 540 ± 25 nm). Baseline was measured for 2-3 min and cells were then challenged with BAM8-22 or vehicle. The BRET signal was calculated as the ratio of the acceptor (YFP or Venus) emission over the donor (RLuc8, Nanoluc) emission. ΔBRET represents the BRET signal in the presence of agonist, minus the mean from 5 initial reads and baseline-corrected to vehicle. Area under the curve (AUC) was determined for each replicate.

### FRET biosensor assays

HEK293 cells were transfected with FRET biosensors (2 µg/10 cm dish), and MrGPRX1 (2 µg/10 cm dish) with or without a co-transfection of Dynamin K44A (1 µg/10 cm dish). After 24 h, cells were transferred to 96-well plates and serum-starved overnight for ERK activity assays. On the day of the assay, cells were washed and incubated in Hanks’ buffered saline solution (HBSS, +10 mM HEPES, pH 7.4) containing 0.1% BSA. In some experiments, cells were pre-incubated in HBSS containing Pitstop 2 or Pitstop 2-negative control (10 µM; Abcam), Dyngo4a (10 µM; Abcam) or Dyngo4a Inactive (10 µM) or vehicle for 30 min, 37°C. FRET was measured at 60 s intervals (CLARIOstar, BMG Labtech). After 5 baseline reads, cells were stimulated with BAM8-22 (1 nM-10 µM). ΔFRET represents the ratio (YFP/CFP), minus the mean from 5 initial reads and baseline-corrected to vehicle. Area under the curve (AUC) was determined for each replicate.

### Luciferase transcriptional assays

HEK293 cells were transfected with luc2P vectors (2 µg/10 cm dish), and MrGPRX1 (2 µg/10 cm dish) with or without a co-transfection of Dynamin K44A (1 µg/10 cm dish).

After 24 h, cells were transferred to 96-well plates and serum-starved overnight. On the day of the assay, cells were treated with vehicle or BAM8-22 (1 µM). After 4 hours (37 °C, 5% CO_2_), cells were incubated with luciferin and lysis buffer for 5 min to enable the substrate reaction. Relative luminescence units (RLU) were measured (CLARIOstar, BMG Labtech).

#### Calcium (Ca^2+^) assays in HEK293 cells

HEK293 cells expressing MrGPRX1, TRPV1, or MrGPRX1-TRPV1 were seeded onto poly-D-lysine coated 96-well clear plates (30,000 cells/well) and cultured for 24 h. Cells were loaded with Fura2-AM ester (1μM) in Calcium Buffer [150 mM NaCl, 2.6 mM KCl, 1.18 mM MgCl2, 2.2 mM CaCl2, 10 mM Glucose, 10 mM HEPES, and 0.5% BSA]c supplemented with probenecid (0.04 mM) and pluronic acid (0.5 μM) for 60 min at 37° C. Fluorescence was measured at 340/380nm excitation and 530 nm emission wavelengths using a FlexStation III plate reader. Baseline measurements were recorded for 45s prior to agonist addition. Cells were incubated for 30-60 min with Go 6983 inhibitor prior to agonist administration. For the sensitization of TRPV1 assay, BAM 8-22 was incubated 5 min prior Capsaicin addition.

### Animals

All animal experiments were adhered to the guidelines recommended by the National Institute of Health, the International Association for the study of Pain, the National Centre for the Replacement, Refinement and Reduction of Animals in Research (ARRIVE) guidelines(60) and were conducted per the Guide for the Care and Use of Laboratory Animals. This study was approved by the Animal Ethics of The New York University Institutional Animal Care and use Committee. Male and female C57BL/6J mice (#00064 Jax®, wild-type) were maintained in a temperature-controlled (22°C) environment with a 12 h light/dark cycle and access to food and water ad libitum. Mice were randomly assigned to experimental groups; group size was based on previous similar studies. Investigators were blind to treatments.

### Collection of mouse tissue

Mice were anesthetized (5% isoflurane) and perfused through the ascending aorta with PBS and then 4% paraformaldehyde in PBS. DRG (L4-L5) were removed, fixed in 4% paraformaldehyde in PBS (1 h, 4°C), cryoprotected in 30% sucrose (24 h, 4°C), and embedded in Optimal Cutting Temperature compound (Tissue Tek). Frozen sections (10-12 μm) were mounted, dried (15 min) and stored (−20°C).

### RNAScope® in situ hybridization

mRNA transcripts were localized in mouse and human tissues using a RNAScopeTM Multiplex Fluorescent Reagent v2 Assay Kit (Advanced Cell Diagnostics Inc.) using the protocol for fresh-frozen tissue as recommended by the manufacturer, except for omission of the initial on-slide fixation step with mouse tissue. Probes to mouse (Mm) Mm-mrgprx1(#488771), and Mm-trpv1 (#313331-C3) were used. To detect all neurons in mouse tissues, hybridized slides were blocked and incubated with guinea pig anti-NeuN antibody (1:500, EMD Millipore, N90) overnight, 4°C. Slides were washed and incubated with goat anti-guinea pig Alexa Fluor® 647 (1:1000) 1 hour, RT. Alternatively, neurons were detected using Nissl staining. Slides were washed, incubated with DAPI and mounted. Sections were imaged using a Leica SP8 confocal microscope with HCX PL APO 40x (NA 1.30) oil objective. The percentage of hybridized positive neurons were quantified and normalized by the total number of neurons.

### Mouse DRG cultures

C57BL/6 mice (∼4 weeks) were euthanized following the guidelines of the American Veterinary Medical Association for the euthanasia of animals. Thoracic and lumbar DRG were excised and dissociated with papain in Hank’s balanced salt solution (HBSS) (60 U/ml, cat# P3125, Millipore Sigma, St. Louis, MO) for 30 min at 37°C, then in collagenase in HBSS (1 mg/ml, cat# C6885, Millipore Sigma) for another 30 min at 37°C under gentle agitation. They were triturated with 1 ml pipette and passed through a 70-μm filter. Dissociated cells were pelleted and resuspended in DMEM containing 10% FBS and 1% penicillin-streptomycin. Neurons were plated on glass coverslips or on 35 mm glass bottom dishes (MatTek, Ashland MA) pretreated with poly-D-lysine (0.1 mg/ml). For AAV NucEKAR transduction, 1 μl of AAV virus was added to each coverslip or dish. Neurons were used within 72 h.

#### Calcium (Ca^2+^) imaging in mouse DRG neurons

Mouse DRG neurons were seeded onto poly-D-lysine coated 35 mm glass bottom dishes (MatTek, Ashland MA) and cultured for 24 h. Changes in agonist-induced calcium influx were determined by loading the neurons with Fura2-AM ester (1μM) in Calcium Buffer [150 mM NaCl, 2.6 mM KCl, 1.18 mM MgCl2, 2.2 mM CaCl2, 10 mM Glucose, 10 mM HEPES, and 0.5% BSA] supplemented with probenecid (0.04 mM) and pluronic acid (0.5 μM) for 60 min at 37 °C. Fluorescence was measured at 340/380nm excitation and 530 nm emission wavelengths using Leica DMi8 Widefield microscope.

Images were taken every ∼10 seconds during the time course of the experiment to minimize photobleaching and phototoxicity and to provide acceptable image quality. Changes in [iCa^2+^] were monitored following a ratio of F340/F380, calculated after subtracting the background from both channels. Baseline measurements were recorded for 60s prior to agonist addition. Cells were incubated for 30-60 min with Go 6983, U0126, Pitstop 2, or Dyngo4a inhibitors prior to agonist administration. Cells were incubated overnight with ERK inhibitor FR 180204 prior to agonist administration. For the sensitization of TRPV1 assay, BAM 8-22 was incubated 10 min prior Capsaicin addition.

### Behavioral test

Dyngo4A, U0126 (ERK inhibitor) and GF109203X (PKC inhibitor) were dissolved in dimethyl sulfoxide (DMSO) into a stock solution and diluted in PBS prior to use.

Dyngo4A (30µM in 10 μL PBS), U0126 (0.5 µg in 5 μL PBS) and GF109203X (0.25 µg in 5 µL PBS) were administered intrathecally with a 30-gauge needle at the cervical level to anesthetized mice, 30 min prior to pruritogen injection. Control mice received an intrathecal injection of the same volume of 1% DMSO in PBS. After each intrathecal injection, 10 µL of 10 µM BAM8-22 in saline was injected intradermally into the shaved nape of each mouse.

Immediately following the BAM8-22 injection, itch was assessed using an automatic and unbiased behavioral spectrometer (Behavior Sequencer, Behavioral Instruments, NJ; BiObserve, DE) (31, 61, 62). The spectrometer comprised a 40-cm^2^ arena with a CCD camera mounted in the center of the ceiling and a door aperture in the front area of the arena. Movement was assessed by a floor mounted vibration sensor and 32 wall mounted infrared transmitter and receiver pairs. Mice were individually placed in the center of the behavioral spectrometer, and their behavior was recorded for 30 minutes and analyzed using a combination of video tracking analysis (Viewer3, BiObserve, Sankt Augustin, Germany) and vibration analysis. Total distance traveled in the open field (track length; m), time still (minutes), and time engaged in grooming (minutes), were recorded and analyzed as described. Scratching behavior was analyzed by the experimenter offline after the record. Each scratching bout was defined as rapid brushing of the back nape by the hind paw.

## Data analysis

Data presented as mean ± SEM. Differences were assessed using paired or unpaired t-test for two comparisons, and 1- or 2-way ANOVA and Tukey’s, Dunnett’s or Šidák’s post-hoc test for multiple comparisons. P<0.05 was considered significant at the 95% confidence level.

## Data availability

All data are contained within the article.

## Conflict of interest

The authors declare that they have no conflicts of interest with the contents of this article.

## Acknowledgments

Author contributions

D.D.J. developed the concept, designed experiments, and supervised all aspects of this project; P.D., J.S.R., and D.D.J. wrote the manuscript. P.D., J.S.R., M.A.F., and K.T. collected and analyzed the data; P.D., J.S.R., M.A.F. and K.T. performed the biochemistry experiments; J.S.R., and M.A.F. performed behavior studies. All authors had the opportunity to discuss the results and comment on the manuscript.

## Funding and additional information

Supported by NIH awards: NIH R01NS125413-01 awarded to DDJ. All data is available in the main text, figures, supplementary materials, and dataset.

## Abbreviations and nomenclature

GPCR (G-protein–coupled receptor), BRET (bioluminescence resonance energy transfer), HEK293 (human embryonic kidney 293 cell line), PEI (polyethyleneimine), MrGPRX1 (Mas-related G-protein coupled receptor X1), TRPV (Transient Receptor Potential Vanilloid)

---

**Supporting Figure 1.**
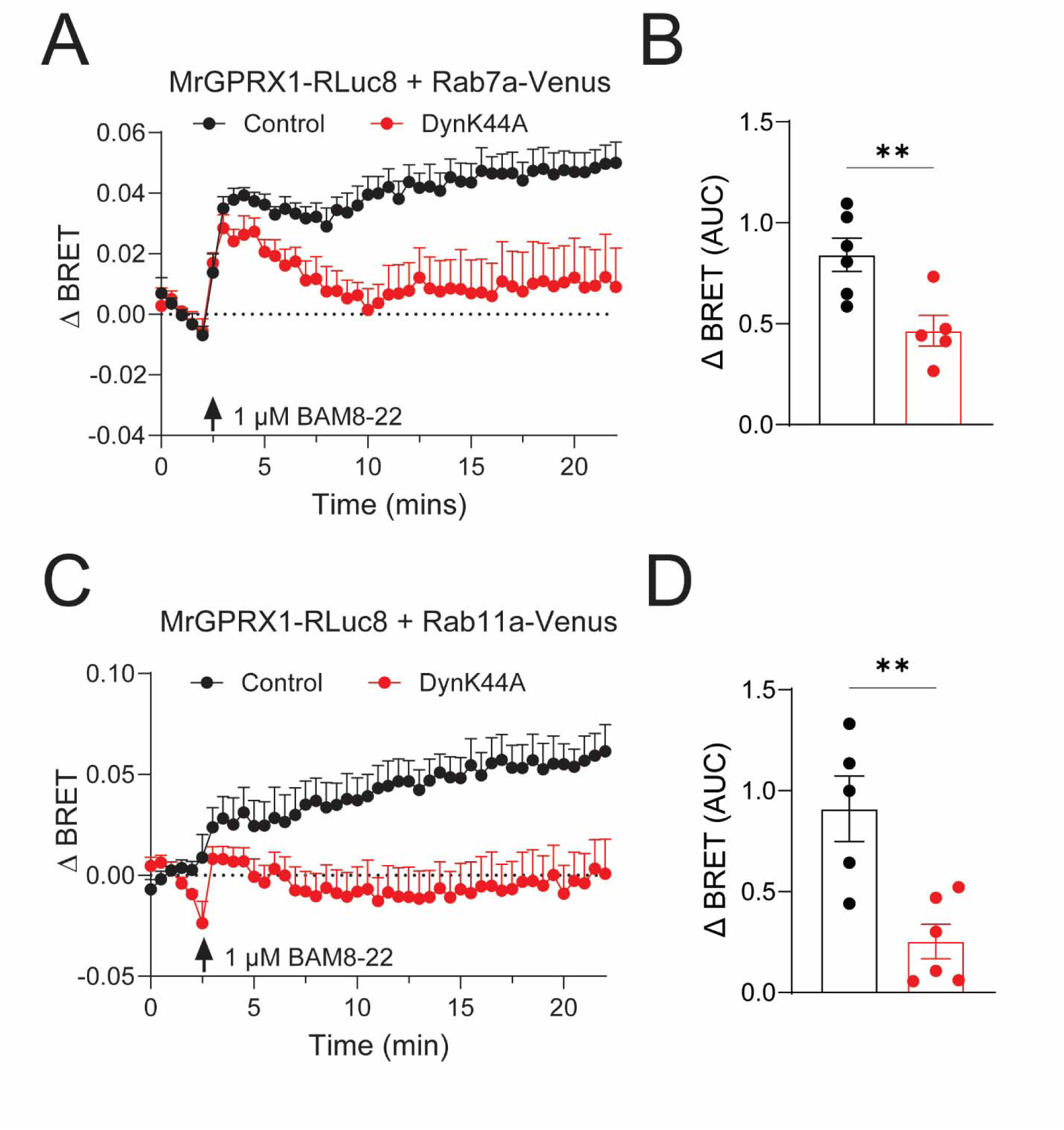
MrGPRX1 traffic can be blocked by dynamin inhibition. BAM8-22-induced (1 µM) trafficking of MrGPRX1-Rluc8 to late endosomes (A) (Rab7a-Venus), and to (C) recycling endosomes (Rab11a-Venus) in HEK293 cells is blocked by DynK44A expression. Area under the curve (AUC) represents quantification of change in BRET over time. n_≥_5, ** P_≤_ 0.01. Unpaired t-test.

**Supporting Figure 2.**
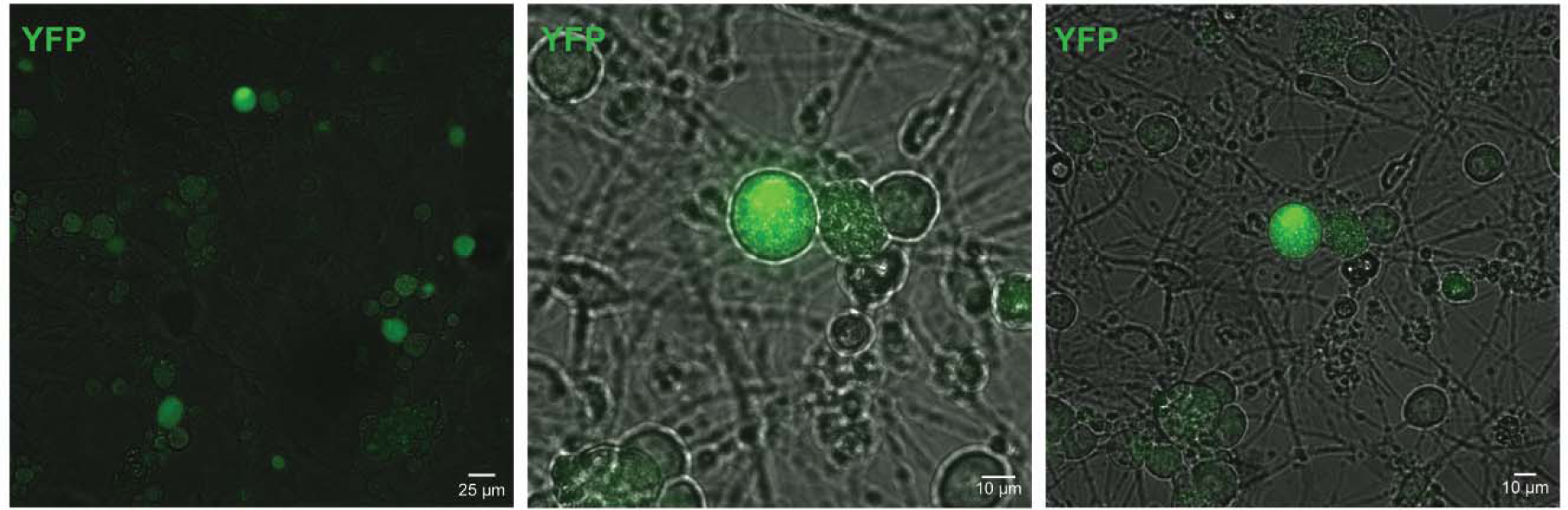
NucEKAR FRET biosensor expression in mouse dorsal root ganglion (DRG) neurons. Representative image of mouse DRG neurons in culture transduced with adeno-associated virus (AAV-NucEKAR) nuclear ERK FRET biosensor. Scale, 25 µm, and 10 µm.

**Supporting Figure 3.**
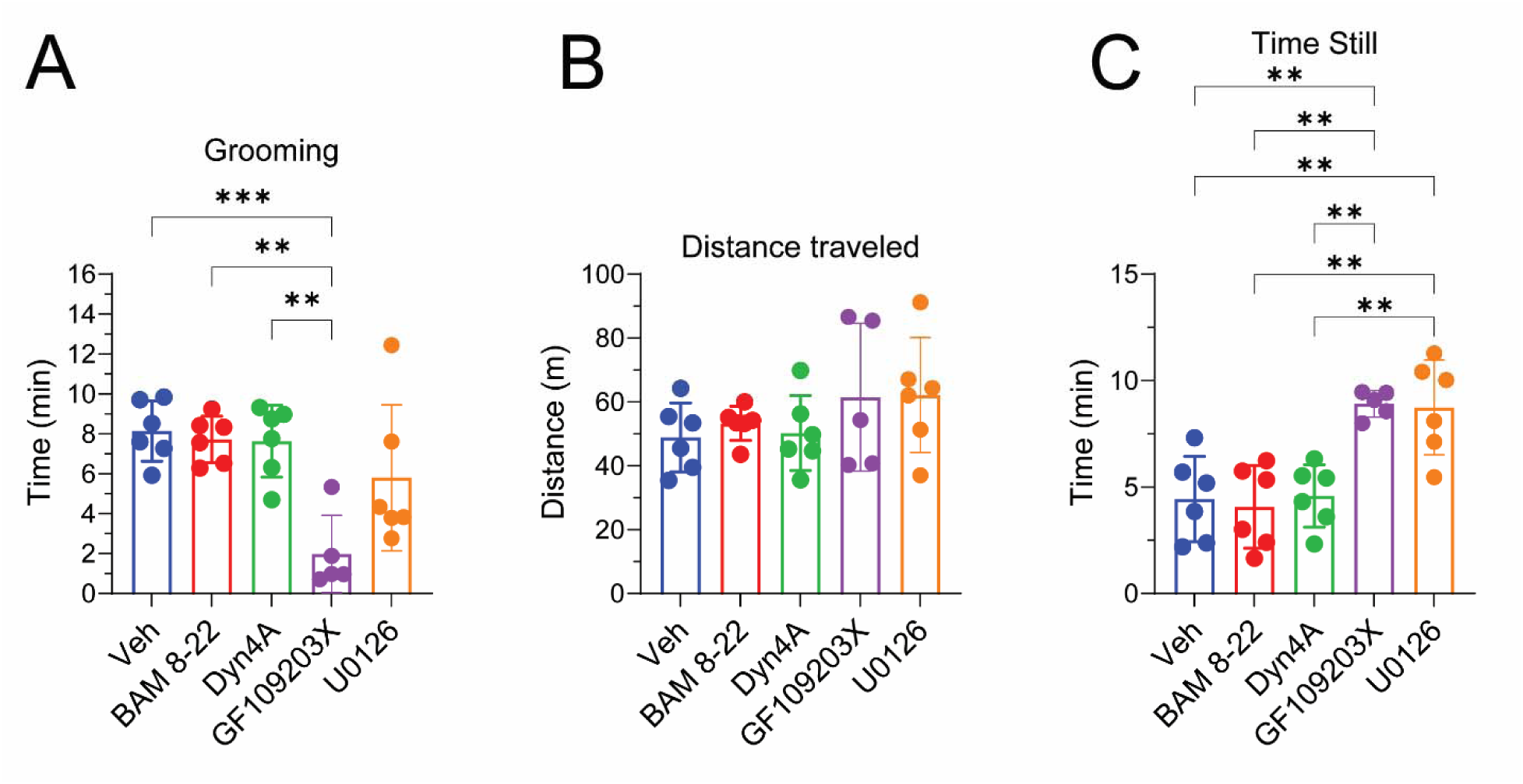
Effect of endocytic, MEK and PKC inhibitors in non-evoked behaviors. Non-evoked behaviors (A) grooming, (B) distance traveled, and (C) time still of vehicle or BAM8-22 intradermal injection 30 min after intrathecal administration of endocytic inhibitor Dyngo4A, MEK inhibitor U0126, or PKC inhibitor GF109203X. n>5 mice, ** P_≤_ 0.01, *** P_≤_ 0.001. One-Way ANOVA, Tukey multiple comparisons.

## Notes

### Competing Interest Statement

The authors have declared no competing interest.

